## Supplemental material for "Drying enhances signal intensity for global GC-MS metabolomics"

**content:**

1. **Supplementary figures**


**Supplementary figure S1**

*Metabolite signal enhancement using drying method.*

**Supplementary figure S2**

*By-products of derivatization reaction decrease using drying method.*

**Supplementary figure S3**

*Drying enhances metabolite signals across different metabolome samples*.

**Supplementary figure S4**

*Drying enhances metabolite signal in human urine samples*.

**Supplementary figure S5**

*Increasing complexity of metabolite mixture is influencing individual metabolite signals*.

**Supplementary figure S6**

*Metabolite signal effect including a drying step in MSTFA derivatization*

**Supplementary figure S7**

*Metabolite signal effect including a drying step in MTBSTFA derivatization*

**Supplementary figure S8**

*3D printed drying device for 24 vials*.

**figure S1**.

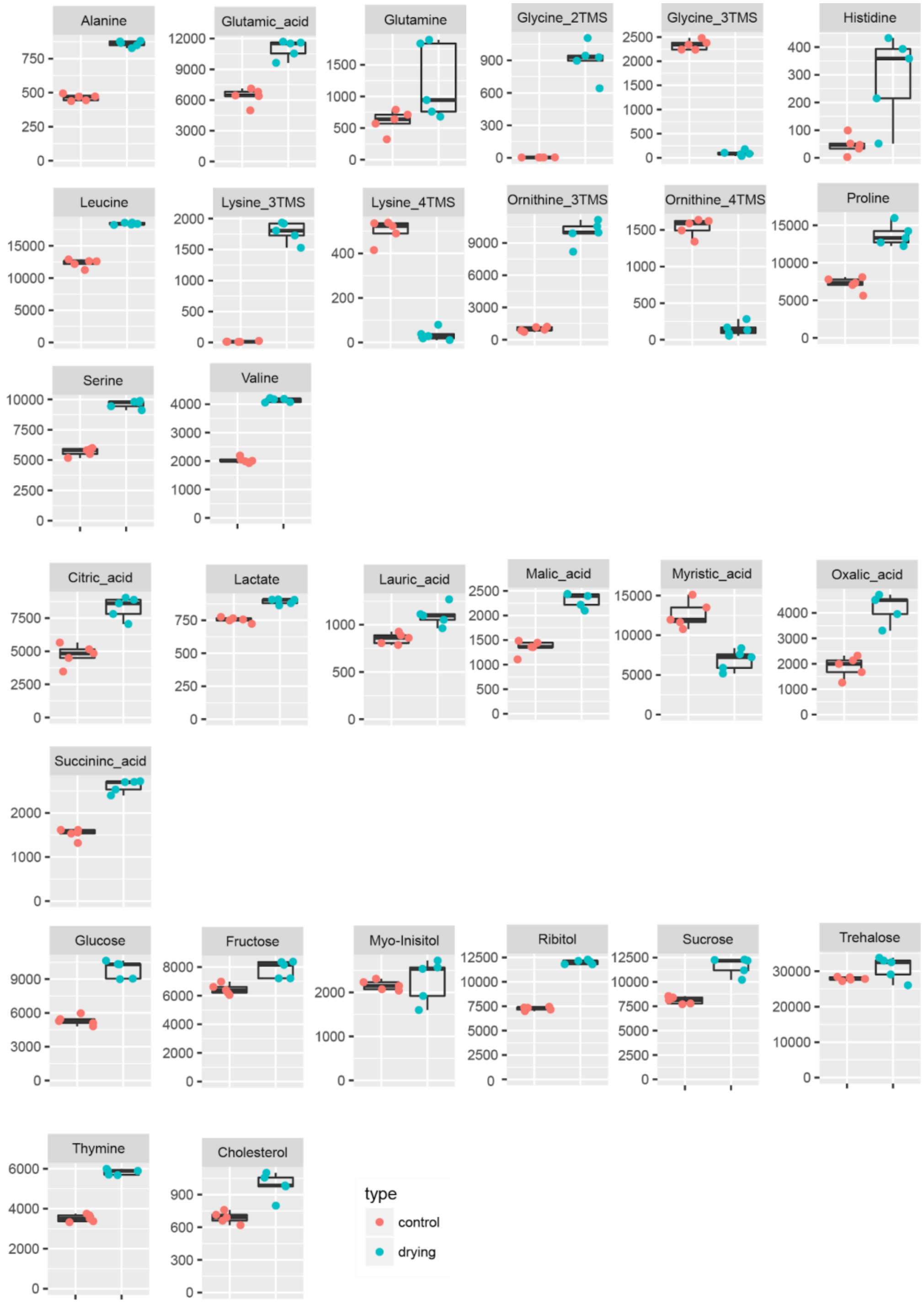


**Supplementary figure S1**. *Metabolite signal enhancement using drying method.* Boxplot graph of all metabolites and their main TMS derivatives from a synthetic metabolite mixture, comparing standard derivatization 'control' vs. derivatization method including drying the methoxymation step 'drying' (each method n=5).

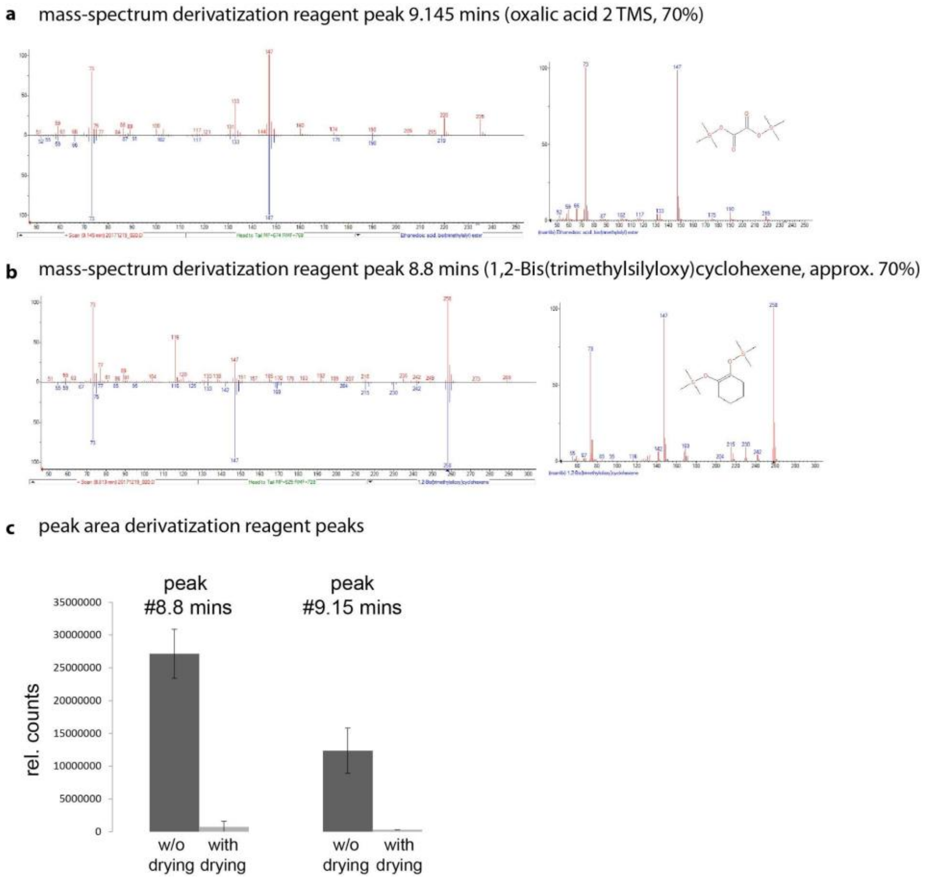


**Supplementary figure S2**. *By-products of derivatization reaction decreased using drying method.* **A)** and **B)** NIST database matches of spectra from peaks with decreased signal intensity using the drying method vs. standard method. **C)** Relative quantification of peaks at 8.8 mins and 9.15 mins analyzed by the drying method vs. standard method. Data from synthetic metabolite mixture, n= 5.

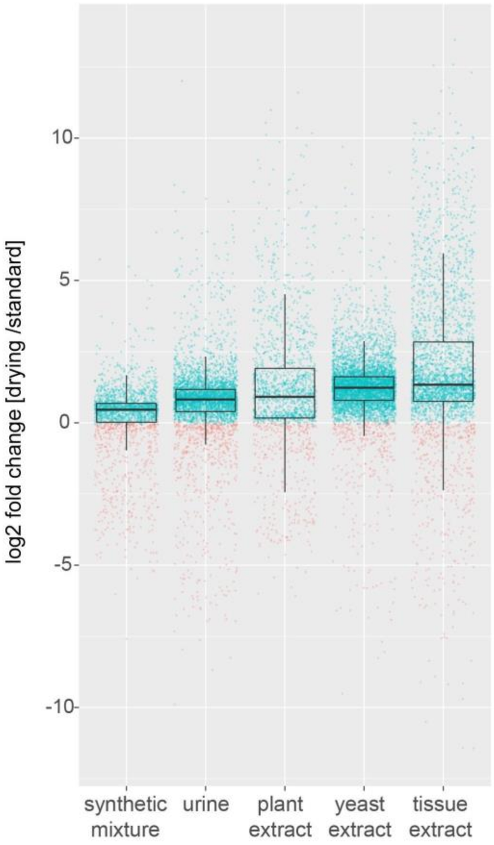


**Supplementary figure S3.** *Drying enhances metabolite signals across different metabolome samples*. Fold change of detected GC-MS features in each sample type, comparing the standard method and the drying method (positive fold change in green, negative fold change in red). Each sample type was analyzed in triplicates using both methods.

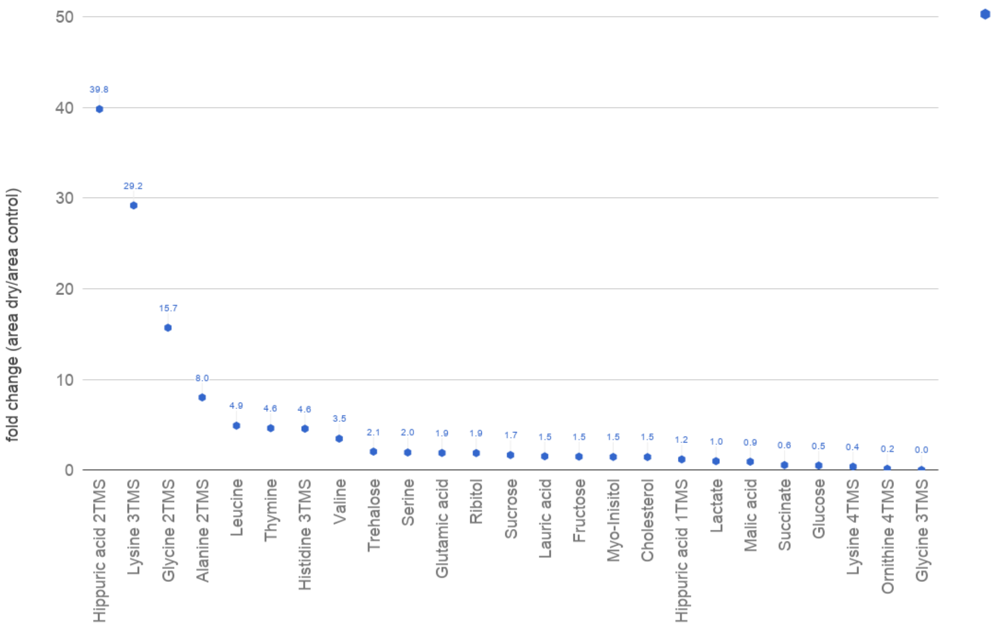


**Supplementary figure S4.** *Drying enhances metabolite signal in human urine samples*. Selected identified metabolites in urine shown as fold-change comparing drying vs. control derivatization method.

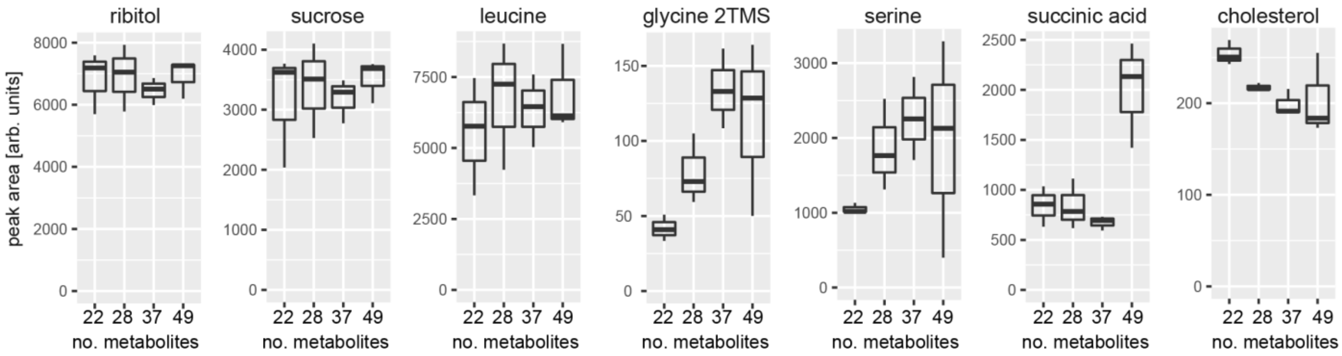


**Supplementary figure S5.** *Increasing complexity of metabolite mixture is influencing individual metabolite signals*. Starting with 22 metabolites, the complexity of the analyzed metabolite mixture was increased in different steps by adding consecutively 6 metabolites, 9 metabolites and 12 metabolites to reach in total of 49 different compounds. Each complexity step increase was done in triplicate.

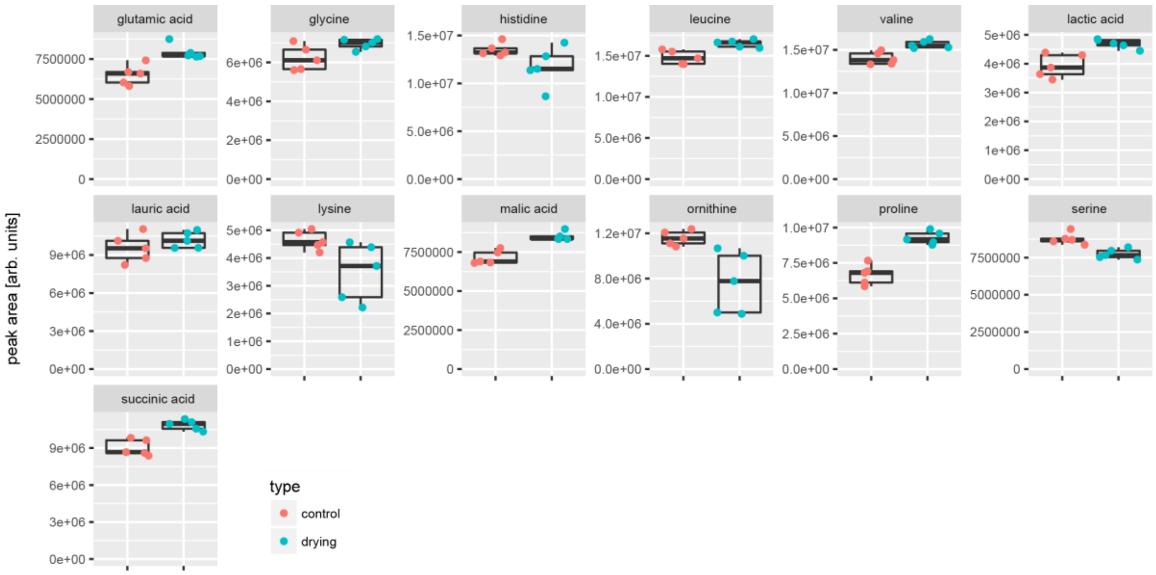


**Supplementary figure 6**. *Metabolite signal effect including a drying step in MTBSTFA derivatization.* Boxplot graph of all major metabolites as their respective t-butyl-dimethylsilyl derivative from a synthetic metabolite mixture, comparing standard derivatization ‘control’ vs. derivatization method including drying the methoxymation step ‘drying’ (each method n=5).

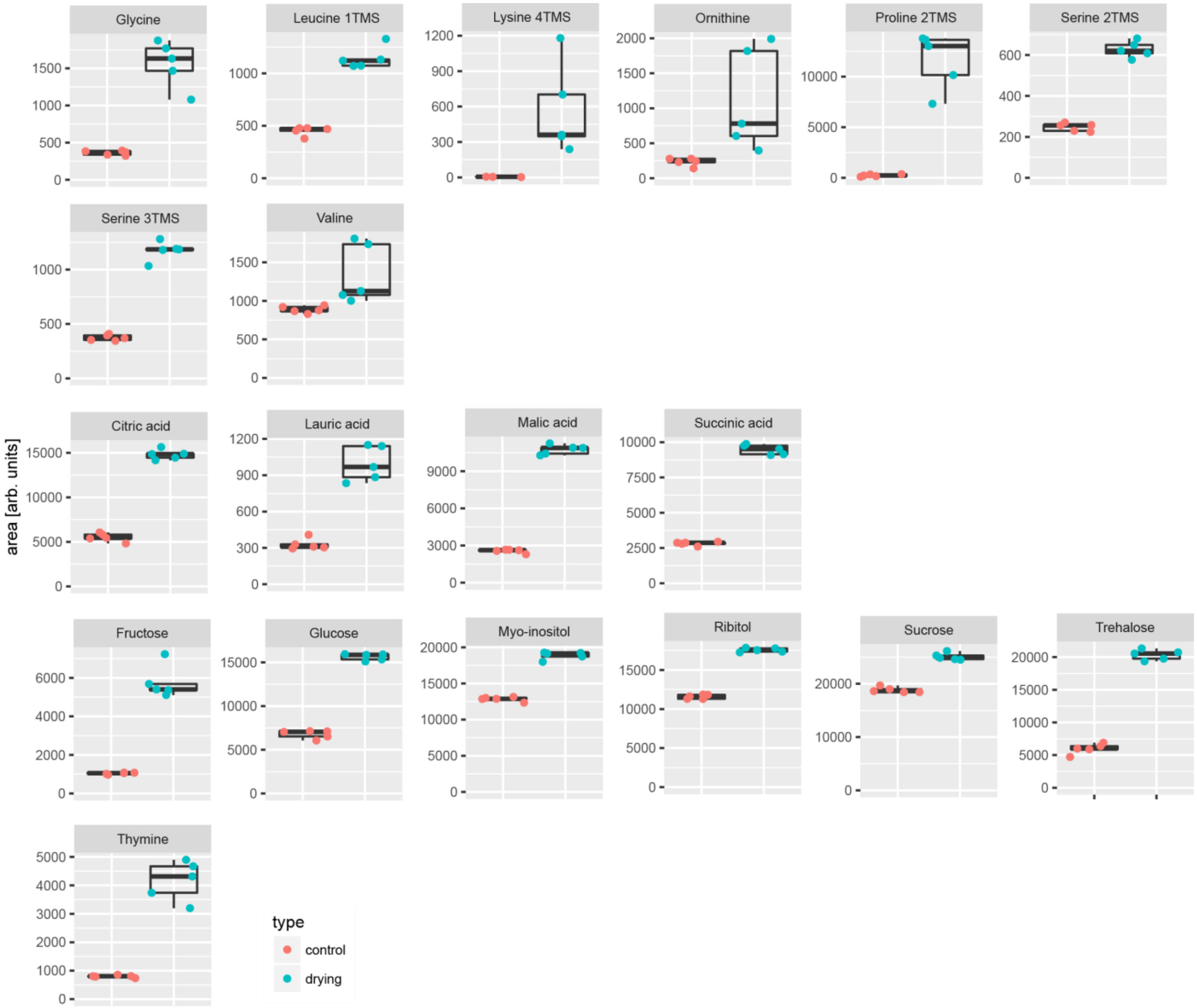


**Supplementary figure S7**. *Metabolite signal effect including a drying step in MSTFA derivatization.* Boxplot graph of all major metabolites as their respective TMS derivative from a synthetic metabolite mixture, comparing standard derivatization ‘control’ vs. derivatization method including drying the methoxymation step ‘drying’ (each method n=5).

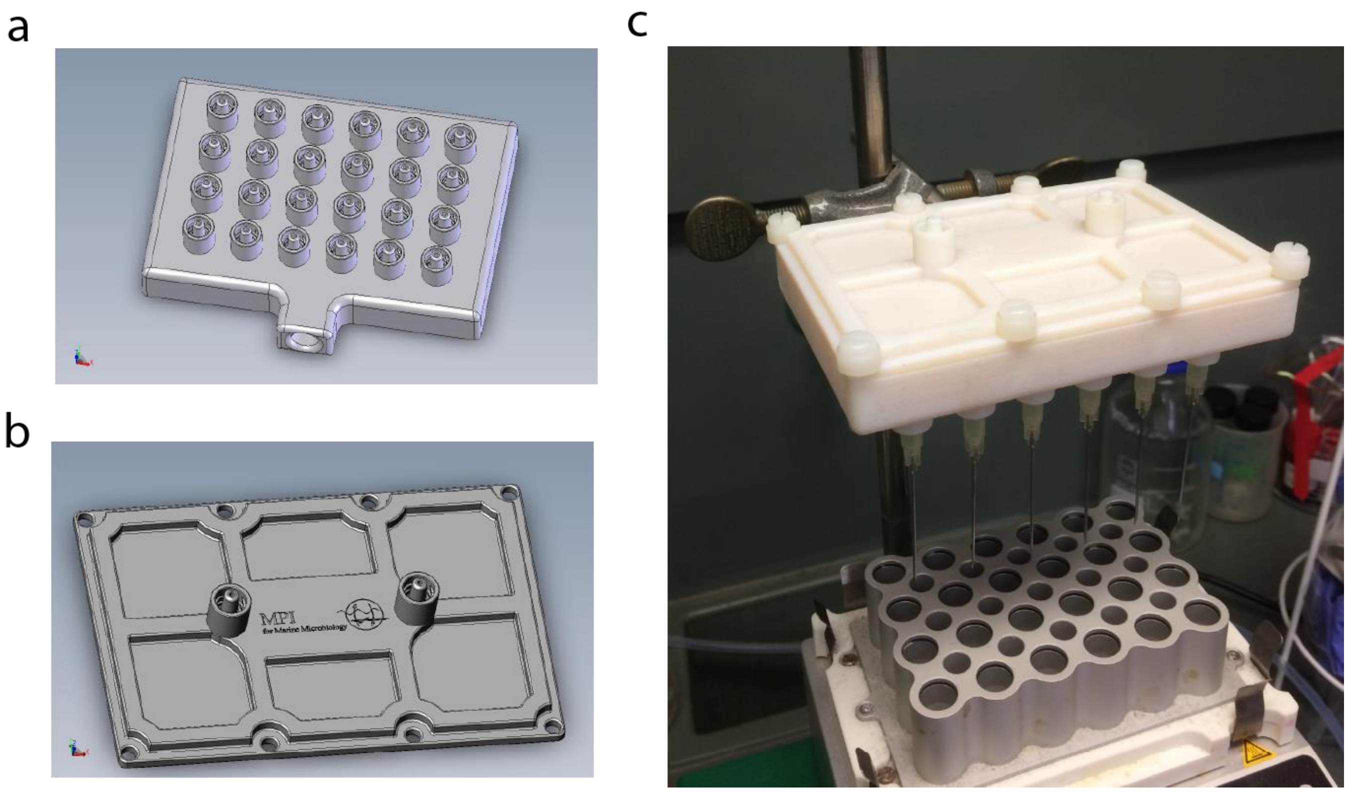


**Supplementary figure S8.** *3D printed drying device for 24 vials*. Model for lower part of device **a)** and upper part to connect nitrogen gas **b)**. **c)** Drying device in the lab, printed with a Stratasys Eden 350V 3D printer and high temperature acrylate based material (RGD 525). The 24 outlets are designed with standard Luer-Lock male connectors to enable the connection of disposable needles. Files can be found (https://figshare.com/s/dd2d8a117d152c8b1fe2).
